# Global broadcasting of local fractal fluctuations in a bodywide distributed system supports perception via effortful touch

**DOI:** 10.1101/2019.12.15.876961

**Authors:** Madhur Mangalam, Nicole S. Carver, Damian G. Kelty-Stephen

**Affiliations:** Department of Physical Therapy, Movement and Rehabilitation Sciences, Northeastern University, Boston, Massachusetts, United States of America; Department of Psychology, University of Cincinnati, Cincinnati, Ohio, United States of America; Department of Psychology, Grinnell College, Grinnell, Iowa, United States of America

**Keywords:** biotensegrity, center of pressure, dynamic touch, effortful touch, multifractality, postural sway, proprioception, psychophysics, tensegrity

## Abstract

A long history of research has pointed to the importance of fractal fluctuations in physiology, but so far, the physiological evidence of fractal fluctuations has been piecemeal and without clues to bodywide integration. What remains unknown is how fractal fluctuations might interact across the body and how those interactions might support the coordination of goal-directed behaviors. We demonstrate that a complex interplay of fractality in mechanical fluctuations across the body supports a more accurate perception of heaviness and length of occluded handheld objects via effortful touch in blindfolded individuals. For a given participant, the flow of fractal fluctuation through the body indexes the flow of perceptual information used to derive perceptual judgments. These patterns in the waxing and waning of fluctuations across disparate anatomical locations provide novel insights into how the high-dimensional flux of mechanotransduction is compressed into low-dimensional perceptual information specifying properties of hefted occluded objects.

## INTRODUCTION

Our smooth perceptuomotor functioning rests on the hardly noticed and rarely studied capability of effortful touch. Our eyes can only face one way, and effortful touch picks up the remaining surroundings. Effortful touch includes perceiving the body, attachments to the body, and the surfaces and substances adjacent to the body. Effortful touch serves as the chief perceptual faculty to the blind when using a cane, or to the sighted when extending a foot forward without looking down or exploring objects just out of view. Effortful touch allows perceiving an intended property of an object (e.g., heaviness, length, width, and shape, orientation in hand) by using various anatomical components (*1*–*9*), or in coordination with each other (e.g., hefting an object using the right vs. left hand, hand vs. foot) (*10*–*16*). In fact, despite the apparent separability of all the disparate anatomical components that can touch, the emerging truth is that no particular anatomical component supports effortful touch in isolation—the arm supports the hand, the torso supports the arm, and the legs support the torso. In this study, we show using causal network modeling that length and heaviness perception of handheld objects via effortful touch in blindfolded humans depends on a complex interplay of mechanical fluctuations across the body.

The neurophysiology subserving effortful touch spans a vast and complex network of connective tissues and extracellular matrix (ECM) that orchestrates the coordination of sensorimotor activity (*17, 18*). Connective tissues distribute tensions and compressions across a wide range of scales and around all parts of the body; this distribution of tension and compression translates local mechanical disturbances into the global realignment of forces (*19*–*23*). Perception via effortful touch emerges from the complex interactions across scales. Specifically, movements during effortful exploration shape the patterns of stimulation available to the body, and the multi-scaled aspect of movement supports a multi-scaled capacity for the body to pick up a wide range of stimulus information, from coarse to fine (*24, 25*). If perception via effortful touch rests on a foundation of action, then it should emerge from the cross-scale interactions of the movement system.

Modeling this connective-tissue support for effortful touch requires a suitable analytical framework. This capacity of the movement system to exhibit across-scale interactions suggests that the bodywide haptic perceptual system may at least exhibit, and at most, depend on, coordination dynamics with the fractal organization (*17, 26, 27*). Indeed, recent work suggests that fractal fluctuations of exploratory movements may have a role in predicting perception via effortful touch. Initial work focused on manual exploration of grasped objects: fractal fluctuations in hand movements improved modeled predictions of perceptual judgments of object properties (heaviness and length) derived by manual hefting (*28*–*30*). Later work investigated the role of postural sway in exploing properties of objects passively supported by the shoulders: just as fractal fluctuations in hand movements had helped predicting perceived properties of manually wielded objects, fractal fluctuations in postural sway also helped predicting perceived properties of objects passively supported by the shoulders (*31, 32*). Besides appearing at multiple contact points between body and perceived object, the predictive role of fractal fluctuations appears to extend across the body: when people are asked to manually heft a grasped object, the relatively distant measure of postural sway, measured as the center of pressure (CoP), has a fractal signature that helps predict the perceptual judgment following hefting (*33, 34*). This cross-body predictive role for CoP fractality increases across trials, indicating progressive implication of fractal fluctuations in perception. Hence, fractal fluctuations provide a window into how specific patterns of movements support specific perceptual goals.

This fractal-shaped window may reveal a coordination of these patterns across the body. It is, of course, possible that CoP fractality is a downstream echo of exploratory patterns at the hand. But an intriguing possibility is that CoP fractality might somehow rise to meet the hand. Specifically, examining how fractality spreads from one distinct anatomical component to another may predict how well these components integrate information supporting the perceptual responses. Indeed, charting out such a relationship has already bore predictive fruit: the effect of visual feedback on judgments via effortful touch depends on fractal fluctuations in head sway as people actively look out on the visible scene, and increases in fractal fluctuations in head sway boost the degree of fractality at the hand (*35*).

The present work aims to tackle the relationship that COP fractality shows with the rest of the body. It specifically answers the following questions: How does the global broadcasting of CoP fractality in a bodywide haptic perceptual system support perception of object properties via effortful touch? For instance, does COP fractality spread upward to the arm? Or is COP fractality spread just the downstream consequence of hefting by the arm? Do the bodywide relationships supporting bodywide flow of fractal fluctuations support more accurate perceptual judgments?

In this study, we investigated how the bodywide dispersal and global broadcasting of local fractal fluctuations across various anatomical locations supports the effortful perception of object properties by manual hefting. We used causal network modeling via vector autoregressive (VAR) analysis (*36*) to capture linear interdependencies among the time series of mechanical fluctuations across multiple anatomical locations to identify the causal network structure of the bodywide haptic perceptual system of effortful touch. So, specifically, we included a set of 13 locations on the body and hefted object during manual exploration to derive perception of heaviness and length, and we tested all possible pairwise relationships between these locations for an exchange of fractal fluctuations. We expected that the waxing and waning of fluctuations across various anatomical locations would provide insights into how bodywide coordinations supported effortful touch. Specifically, we predicted both that CoP fractality would promote fractal patterning in the arm and that the strength of statistically significant pairwise exchanges of fractal fluctuations would serve to predict greater accuracy (i.e., lower absolute errors).

## RESULTS

### Hefting objects to perceive heaviness and length

Fifteen blindfolded healthy adults hefted with their right hand six specially-designed experimental objects that systematically differed in their their mass, *m* (Object 1 > Object 2, Object 3 > Object 4, Object 5 > Object 6), the static moment, **M** (Object 1 = Object 2 = **M**_S_ < Object 3 = Object 4 = **M**_M_ < Object 5 = Object 6 = **M**_L_), and the moment of inertia, *I*_1_ and *I*_3_, reflecting the resistance of the object to rotation about the longitudinal axis (*I*_1_ values: Object 1, Object 2, Object 3 < Object 4, Object 5 < Object 6) (Table 1 and Fig. 1A). To introduce variability in manual exploration, we introduced anatomical and kinematic constraints on manual exploration. The participants hefted each object as their wrist was constrained to move about 10° radial deviation (Fig. 1C, top panels), the neutral position (Fig. 1C, middle panels), or 10° radial deviation (Fig. 1C, bottom panels). In a static condition, the participant lifted and held each object static (Fig. 1C, left panels). In two separate dynamic conditions, the participant lifted and wielded each object synchronously with metronome beats at 2 Hz or 3 Hz (Fig. 1C, center and right panels, respectively). The participant assigned heaviness values proportionally higher and lower than 100 to an object perceived heavier and lighter, respectively, than the reference object (e.g., 200 to an object perceived twice as heavy and 50 to an object perceived half as heavy). They reported perceived length of the object by adjusting the position of a marker along a custom string-pulley assembly.

**Table 1.**
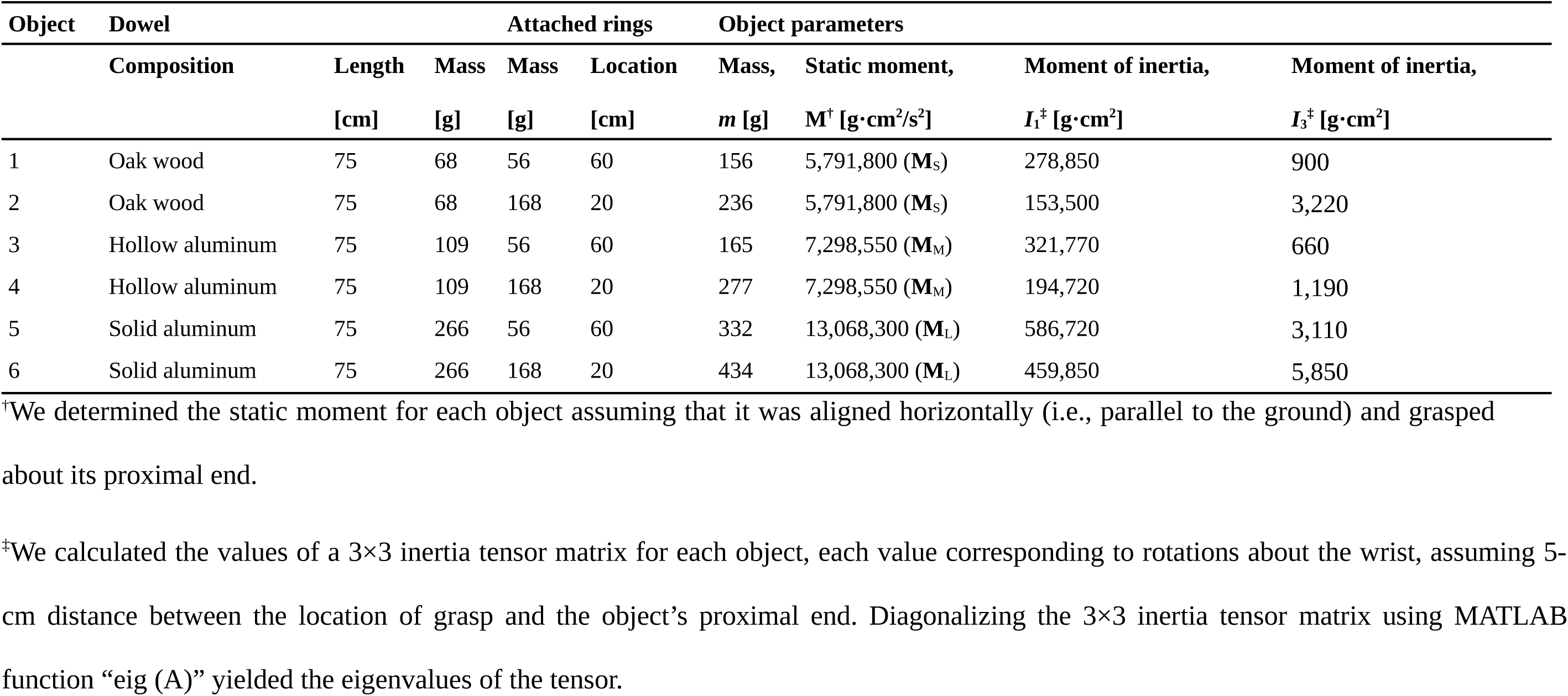
Experimental objects.

| Object | Dowel |  | Attached rings |  |  | Object parameters |  |  |  |
| --- | --- | --- | --- | --- | --- | --- | --- | --- | --- |
|  | Composition | Length | Mass | Mass | Location | Mass, | Static moment, | Moment of inertia, | Moment of inertia, |
| | | [cm] | [g] | [g] | [cm] | $m$ [g] | $M^{\dagger}$ [g·cm<sup>2</sup>/s<sup>2</sup>] | $I_1^{\dagger}$ [g·cm<sup>2</sup>] | $I_3^{\dagger}$ [g·cm<sup>2</sup>] |
| 1 | Oak wood | 75 | 68 | 56 | 60 | 156 | 5,791,800 ($\mathbf{M}_S$) | 278,850 | 900 |
| 2 | Oak wood | 75 | 68 | 168 | 20 | 236 | 5,791,800 ($\mathbf{M}_S$) | 153,500 | 3,220 |
| 3 | Hollow aluminum | 75 | 109 | 56 | 60 | 165 | 7,298,550 ($\mathbf{M}_M$) | 321,770 | 660 |
| 4 | Hollow aluminum | 75 | 109 | 168 | 20 | 277 | 7,298,550 ($\mathbf{M}_M$) | 194,720 | 1,190 |
| 5 | Solid aluminum | 75 | 266 | 56 | 60 | 332 | 13,068,300 ($\mathbf{M}_L$) | 586,720 | 3,110 |
| 6 | Solid aluminum | 75 | 266 | 168 | 20 | 434 | 13,068,300 ($\mathbf{M}_L$) | 459,850 | 5,850 |

<sup>†</sup>We determined the static moment for each object assuming that it was aligned horizontally (i.e., parallel to the ground) and grasped about its proximal end.
<sup>‡</sup>We calculated the values of a 3×3 inertia tensor matrix for each object, each value corresponding to rotations about the wrist, assuming 5- cm distance between the location of grasp and the object's proximal end. Diagonalizing the 3×3 inertia tensor matrix using MATLAB function “eig (A)” yielded the eigenvalues of the tensor.

**Fig. 1.**
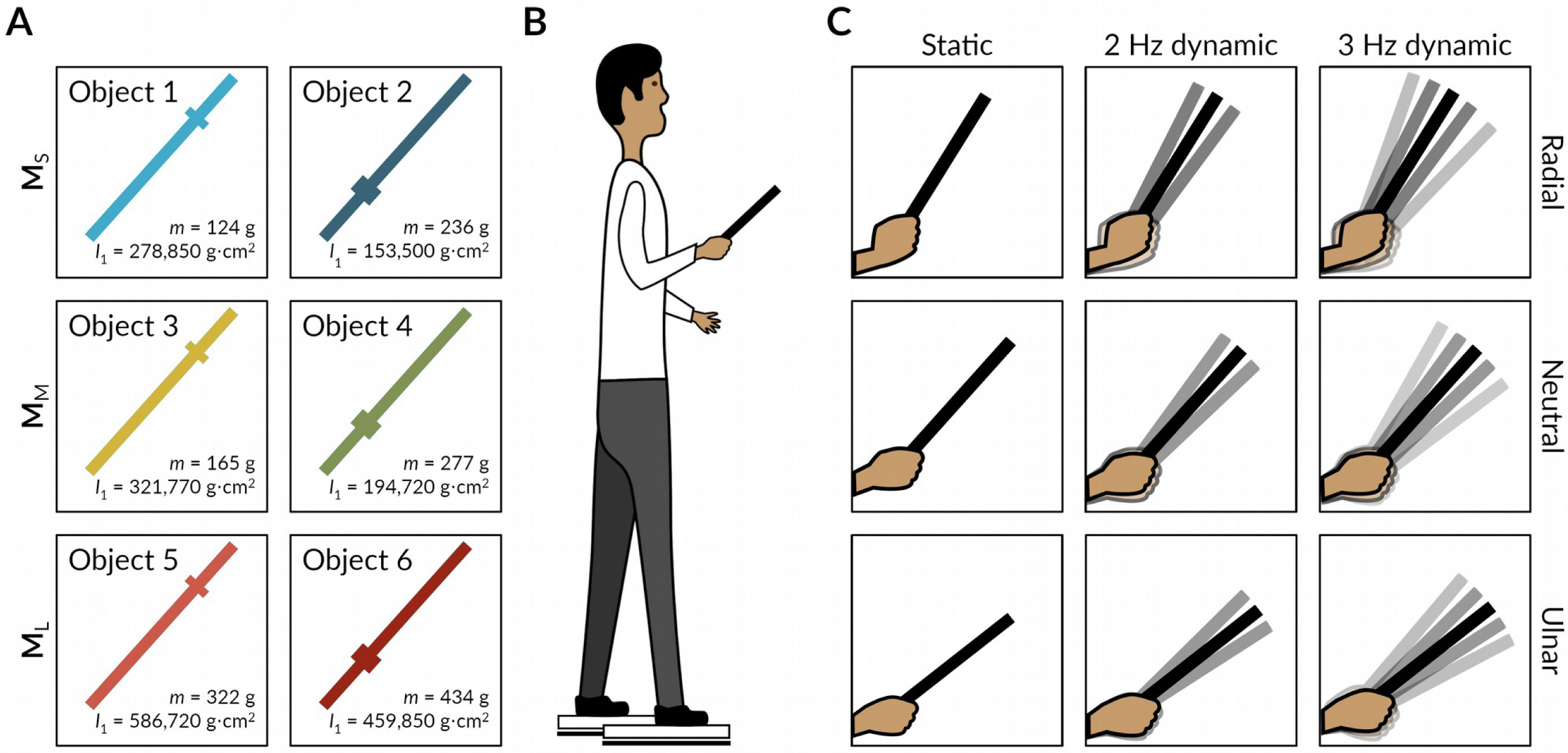
Schematic illustration of the experimental objects, setup, and exploratory conditions. (**A**) Each participant hefted six objects with different mass, m, and the moment of inertia, I_1_. (**B**) Each participant stood with his/her two feet on separate force plates, hefted each object for 5 s, and reported his/her judgments of heaviness and length of that object. (**C**) The participant was instructed to constrain the wrist motion either about 10° radial deviation (top panels), the neutral position (middle panels), or 10° radial deviation (bottom panels). In a static condition (left panels), the participant lifted and held each object static, and in two dynamic conditions, the participant lifted and wielded each object synchronously with metronome beats at 2 Hz or 3 Hz (center and right panels).

### Each anatomical location showed fractal fluctuations

We measured the center of pressure (CoP) and 3D motion of twelve reflective markers attached to the hefted object (*N* = 3) and the participant’s body (n = 9; Supplementary Table S1 and Fig. 2A). Next, we computed a planar Euclidean displacement (PED) series describing fluctuations in CoP between each consecutive sample (Fig. 2B). We also computed a spatial Euclidean displacement (SED) series for each reflective marker describing fluctuations at the respective anatomical location (Fig. 2B).

**Fig. 2.**
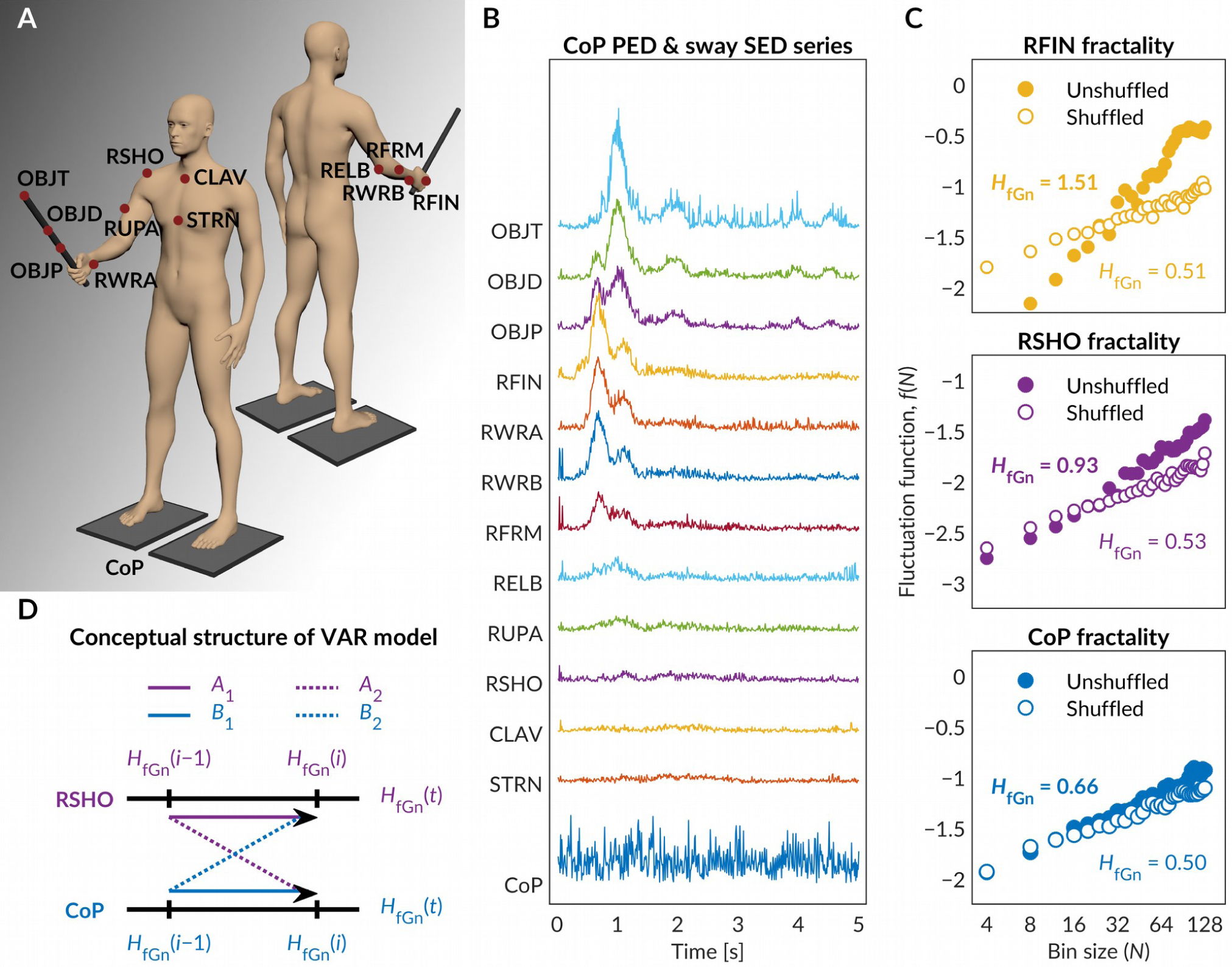
Overview of data acquisition process and analysis. (**A**) Locations of the reflective markers attached to the experimental object and the participant’s body. (**B**) The time series of the planar Euclidean displacement (PED) of CoP and spatial Euclidean displacement (SED) for each of the 12 reflective markers. (**C**) Log-log plots of the fluctuation function, *f*(*N*), vs. bin size reflecting the fractal scaling exponent, *H*_fGn_, yielded by the detrended fluctuation analysis (DFA) in a representative trial. Solid circles and solid trend line describe *f*(*N*) for the original time series; and open circles and dashed trend line describe f(N) for a shuffled version of the original time series. (**D**) The conceptual structure of the vector autoregressive (VAR) analysis used to model the diffusion of fractal fluctuations across different anatomical components. The contribution of each location is represented as a time series of trial-by-trial values of H_fGn_. Arrows represent weights in the model, indicating the effects of fractality in the previous trail on fractality in the current trial.

To test for fractality in CoP PED and each marker SED series, we obtained detrended fluctuation analysis (DFA) estimates of *H*_fGn_ for the original version (i.e., unshuffled) and a shuffled version of each series (Fig. 2C). The random shuffling of a series destroys the temporal structure of a signal, and consequently, any existing temporal correlations characterizing its fractality also disappears. A truly fractal signal yields the fractal scaling exponent *H*_fGn_ > 0.5 as well as *H*_fGn_ greater than the scaling exponent calculated for shuffled series of the same numbers (*37, 38*).

DFA estimates of H_fGn_ for CoP PED series (*Mean* = 0.57, *SEM* = 0.0018) fell in the fractal range (i.e., 0.5 < *H*_fGn_ < 1), and significantly exceeded H_fGn_ for the shuffled versions of the series (*Mean* = 0.51, *SE* = 0.0013), paired-samples *t*-test: *t*_1619_ = 25.57, *P* < 0.001. The same was also true for each marker SED series (all *P*s < 0.001; Supplementary Table S2). Data exploration at the level of individual trials indicated inflection points in fluctuation functions, specifically at larger timescales. We thus tested whether such inflection points may have artificially amplified the values of *H*_fGn_. DFA estimates of *H*_fGn_ for the original version and a shuffled version of the PED series for a shorter, half of the scaling region also yielded similar results (all *P*s < 0.001; Supplementary Table S3), confirming that the inflection points did not artificially amplify the values of *H*_fGn_. Collectively, these results strongly show that fluctuations in CoP and different anatomical locations display fractality.

### Fractality spreads across the body

We used the vector autoregressive (VAR) analysis to model the diffusion of fractal fluctuations among the distinct anatomical components (Fig. 2D). VAR modeling yielded forecasts of the effects of fractality at each anatomical location on fractality at each other anatomical location, as well as at that location itself, in the subsequent ten trials. The dynamic interaction within each possible pair of endogenous variables (i.e., variables that constitute the system itself) were represented by impulse-response functions (IRF) that describes the reaction of one endogenous variable to an impulse in the other variable in the subsequent trials (Fig. 3).

**Fig. 3.**
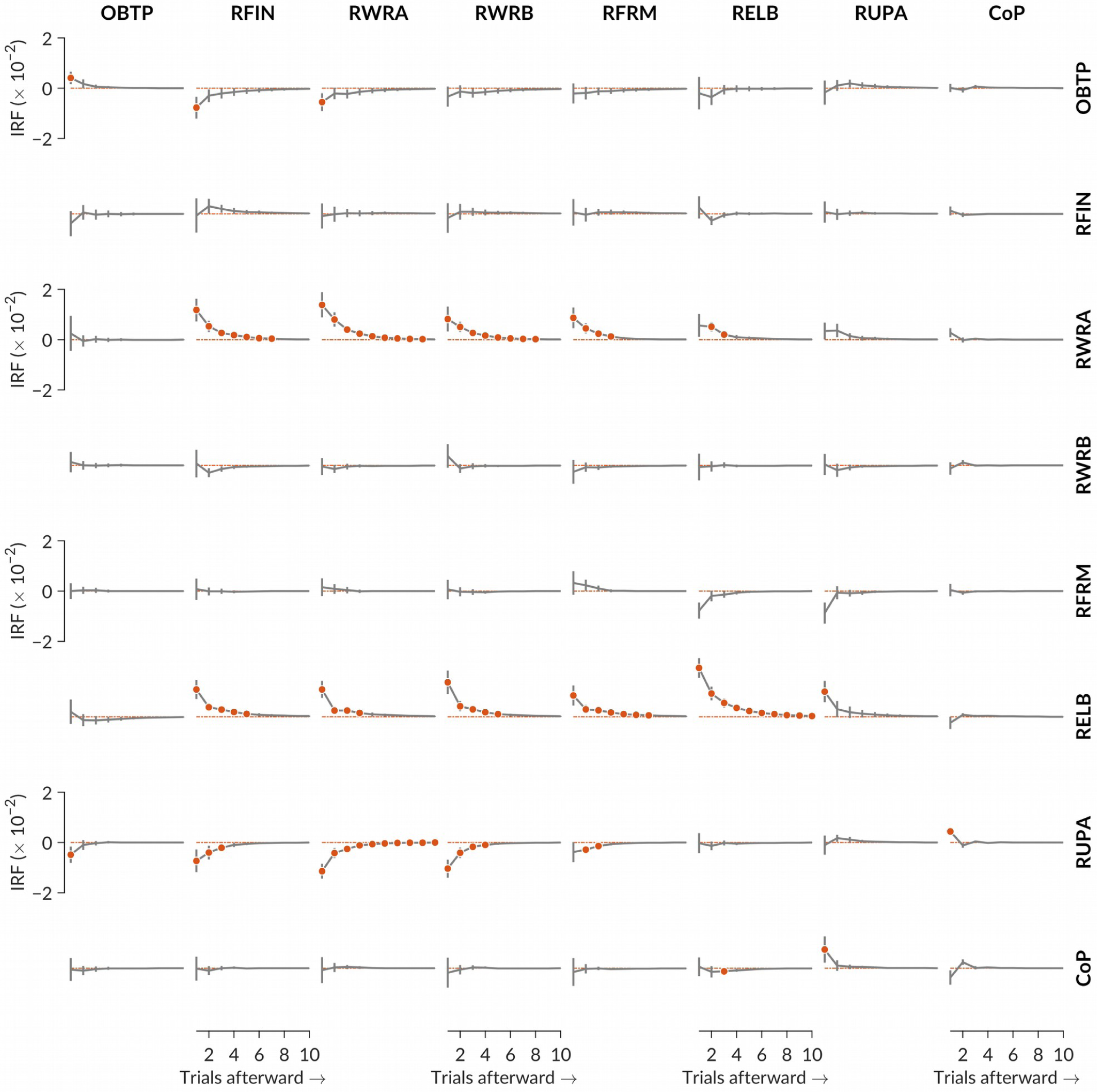
*Mean* (± 1*SEM*) values of impulse-response functions (IRFs) predicting the response of each anatomical component over 10 trials afterward to an impulse in fractality of each other anatomical component in the current trial. For each IRF curve in each panel, row labels indicate impulses, and column labels indicate responses. Each red solid circle indicates a statistically significant (*P* < 0.01) response to an impluse in *i*th trial (1 through 10). Increase in OBTP fractality showed an immediate positive effect on the subsequent values of itself, but this trend diminished fast. Increase in OBTP fractality also showed an immediate negative effect on subsequent fractality of RFIN and RWRA fractality. Increases in RWRA fractality showed a positive effect on subsequent values of RFIN, RWRB, RFRM, and RELB fractality, as well as on subsequent values of itself. Increases in RELB fractality showed a positive effect on subsequent values of RFIN, RWRA, RWRB, RFRM, and RUPA fractality, as well as on subsequent fractality of itself. However, increases in RUPA fractality showed a negative effect on subsequent values of OBTP, RFIN, RWRA, RWRB, and RFRM fractality, suggesting that RUPA increases came at the expense of fractality throughout the arm. Interestingly, RUPA and CoP fractality showed an increasingly positive effect on subsequent fractality of each other. Each of these curves eventually approaches zero, indicating that this effect weakened over subsequent trials and eventually diminished completely.

An increase in OBTP fractality showed an immediate positive effect on the subsequent values of itself, but this trend diminished fast. The object is a simple rigid body without internal degrees of freedom, but the short-range propagation of fractality is an expectable consequence of simple properties like inertia, even for simple systems (*39*). An increase in OBTP fractality also showed an immediate negative effect on subsequent fractality of RFIN and RWRA fractality, suggesting that OBTP fractality’s increases came at the direct expense of finger and wrist fractality. These results make good sense, especially because the object is a passive recipient of fluctuations from the hand, and any fluctuations in the object should be directly the consequence of fluctuations flowing from the arm.

The most distinctive of these IRF relationships suggest that the wrist and elbow facilitated the propagation of fractality through the arm (Fig. 3). An increase in RWRA and RELB fractality promoted subsequent increases in RFIN, RWRB, RFRM fractality, as well as subsequent increases in RWRA and RELB fractality themselves. However, whereas the wrist and elbow were the broadcasters of fractality, it appeared that fractality at the upper arm served to draw fractality away from the arm, as RUPA fractality increased at the expense of RELB fractality.

RUPA fractality appeared to support subsequent increases in COP fractality; and reciprocally, COP fractality appeared to promote subsequent increases in RUPA fractality as well. Our regression modeling confirmed that the individual mean differences from zero, as indicated by the solid red circles in Fig. 3, are, in fact, significant even after controlling for multiple comparisons across all 165 IRF relationships considered (Supplementary Table S4). Hence, fractality from CoP does have consequences for the arm during hefting, but rather than promoting fractality through the rest of the arm, CoP actively drew fractality away. COP fractality promoted later RUPA fractality. However, rather than spreading fractality all the way from RUPA to the rest of the arm, this upwards influence of CoP fractality actually drew down the fractality of the rest of the arm. COP fractality promoted RUPA fractality, leading RUPA to draw fractality from the lower parts of the arm and pass it on towards COP. In short, the wrist and elbow spread fractality to their neighbors. As fractality in the upper arm increased, it brought down fractality among these neighbors (as well as the wrist). Finally, the upper arm and CoP fed upon each other’s fractality. Here we have a potential explanation of how COP fractality in previous work bore the imprint of fractal patterning by manual hefting (*33, 34*).

Thus, hefting an object to perceive the heaviness and length of that object results in a multifarious cascade of effects, spanning across the whole body. The haptic perceptual system benefits from the spreading of fractal fluctuations, thus bearing a close resemblance to complex stochastic networks that exhibit continuous exchange of flows (*40*–*42*). When perturbed, a mechanically organized stochastic network of the kind of the bodywide haptic perceptual system is bound to act to disperse the forces applied to one part of the system to the neighboring parts through ultra-fast diffusion of forces. One corollary of this treatment of the perceptual system is that the perceptual process is not limited to the brain or neurons, and thus clear distinctions between the roles of neural dynamics and bodily mechanics in effortful touch may not be possible. Relinquishing such arbitrary distinctions between neural dynamics and bodily mechanics provides an avenue for novel insights into the functioning of the perceptual system, to which the present findings testify.

### Greater diffusion results in more accurate perception

We hypothesized that if the propagation of fractality across various anatomical locations aids perception, then the individuals who show stronger IRF impulse-responses would show greater accuracy in perceptual judgments. To model the effects of the strength of the propagation of fractality on the accuracy of perception at the individual level, we determined absolute errors in perception of heaviness and length. Because perceived heaviness followed a proportion relative to the reference object of 109-gm, we calculated this judgment as the percentage of the [theoretically] accurate percentage value based on each object’s actual mass. For instance, if a participant attributes to Object 2 (mass = 236 g) a heaviness value of 120 relative to 100 of the referenced object, then they showed an absolute error in perceived heaviness, *H*_error_ = 100 – 100×((120×109)/100)/236 = 44.66. We calculated the absolute error in perceived length, *L*_error_, simply as the absolute values of the difference between the actual length (75 cm) and perceived length.

A generalized linear model (GLM) of Poisson regression revealed that above and beyond that known effects of experimental manipulations, object parameters, and trial order (Table 2) (*33*), the subsequent increase in RFIN fractality due to RELB fractality reduced *H*_error_ (*z* = –1.99, *p* = 0.047; Fig. 4), suggesting that absolute error in perceived heaviness decreased significantly as RELB fractality prompted an increase in RFIN fractality. A linear mixed-effects (LME) model revealed that, above and beyond that known effects of experimental manipulations, object parameters, and trial order (Table 2) (*33*), the subsequent increase in CoP fractality due to RUPA fractality (*t* = –4.00, *P* = 0.007), RWRB fractality due to RFRA fractality (*t* = –3.82, *P* = 0.009), and RWRA fractality due to RELB fractality (*t* = –8.15, *P* < 0.001) reduced *L*_error_ (Fig. 4). At the same time, the flow of fractality from RELB to RWRB increased *L*_error_ (*t* = 4.59, *P* = 0.004; Fig. 4). Hence, most exchanges of fractality across the body supported greater accuracy, except the flow of fractality from the elbow to the wrist.

**Table 2.** Coefficients of generalized linear model (GLM) of Poisson regression and linear mixed-effects (LME) model examining the strength of fractal fluctuations in PED series on the unsigned error in perceived heaviness and perceived length, respectively

|  | Perceived heaviness |  |  | Perceived length |  |  |
| --- | --- | --- | --- | --- | --- | --- |
| **Effects** | ***b* ($\pm 1SEM$)** | ***z*** | ***P*<sup>‡</sup>** | ***b* ($\pm 1SEM$)** | ***t*** | ***P*<sup>‡</sup>** |
| (Intercept) | -51.91 (7.07) | 7.34 | < **0.001** | -205.79 (142.12) | 1.45 | 0.148 |
| $H_{\text{perceived}}$ | | | | 0.0025 (0.0049) | 0.51 | 0.613 |
| Trial order | -0.0092 (0.0036) | 2.60 | **0.009** | 0.029 (0.013) | 2.27 | **0.023** |
| Wrist angle (Radial – Neutral) | 0.044 (0.010) | 4.33 | < **0.001** | -0.26 (0.48) | -0.55 | 0.579 |
| Wrist angle (Ulnar – Neutral) | -0.13 (0.010) | -12.31 | < **0.001** | 1.79 (0.47) | 3.80 | < **0.001** |
| Wrist angular kinematics | -0.00055 (0.0033) | -0.17 | 0.868 | 0.0040 (0.15) | 0.026 | 0.979 |
| Log$I_1$ | 6.27 (1.03) | 6.09 | < **0.001** | 43.81 (25.92) | 1.69 | 0.091 |
| Log$I_3$ | 6.38 (0.70) | 9.13 | < **0.001** | | | |
| as.ordered(**M**).L<sup>†</sup> | -3.03 (0.46) | -6.53 | < **0.001** |  |  |  |
| as.ordered(**M**).Q<sup>†</sup> | -4.14 (0.40) | -10.36 | 0.564 |  |  |  |
| $H_{\text{fGn}}$ at CoP | 119.50 (12.56) | 9.51 | **0.004** | 602.40 (255.11) | 2.36 | 0.018 |
| RFRA on RFIN | -6.82 (4.28) | -1.60 | 0.110 | -221.66 (97.44) | -2.28 | 0.063 |
| RFRA on RWRB | 3.42 (4.11) | 0.83 | 0.406 | -362.87 (94.88) | -3.82 | **0.009** |
| RFRA on RELB | 0.95 (3.11) | 0.30 | 0.761 | 117.05 (70.05) | 1.67 | 0.146 |
| RELB on RFIN | -10.32 (5.20) | -1.99 | **0.047** | -137.82 (132.64) | -1.04 | 0.338 |
| RELB on RWRA | 3.98 (6.05) | 0.66 | 0.511 | -1237.78 (151.94) | -8.15 | < **0.001** |
| RELB on RWRB | 5.87 (4.95) | 1.19 | 0.236 | 582.55 (126.99) | 4.59 | **0.004** |
| RUPA on CoP | -5.35 (7.86) | -0.68 | 0.496 | -798.77 (199.63) | -4.00 | **0.007** |
| CoP on RUPA | -4.27 (2.41) | -1.77 | 0.076 | 119.23 (54.20) | 2.20 | 0.070 |
| Trial order $\times H_{\text{fGn}}$ at CoP | -0.017 (0.0064) | -2.66 | **0.008** | | | |
| $H_{\text{perceived}} \times$ Trial order | | | | -0.00010 (0.000070) | -0.86 | 0.339 |
| $H_{\text{fGn}}$ at CoP × as.ordered(**M**).L<sup>†</sup> | 6.72 (0.83) | 8.12 | < **0001** | | | |
| $H_{fGn}$ at $CoP \times as.ordered(\mathbf{M}).Q^\dagger$ | 7.66 (0.71) | 10.75 | < **0.001** | | | |
| $H_{fGn}$ at $CoP \times LogI_1$ | -15.17 (1.83) | - 8.29 | < **0.001** | -109.82 (46.52) | - 2.36 | **0.018** |
| $H_{fGn}$ at $CoP \times LogI_3$ | -10.93 (1.25) | - 8.76 | < **0.001** | | | |

<sup>†</sup>These listings indicate the default treatment of an ordinal variable. Because the spacing between levels of ordinal variables may not necessarily be even, the best statistical practice for modeling the effect of an ordinal variable with $k$ levels is to fit the orthogonal polynomials of order 1 to $k - 1$ . Accordingly, we included the linear (L) and quadratic (Q) effects of the static moment ( $\mathbf{M}$ ) in the model to control for any nonlinear effect of $\mathbf{M}$ and to test whether $H_{fGn}$ moderates the effect of $\mathbf{M}$ . Importantly, both the linear (L) and quadratic (Q) effects of $\mathbf{M}$ are warranted on statistical grounds to represent the effect of $\mathbf{M}$ accurately, and neither effect is specifically relevant for theoretical reasons. Our theory suggests simply that $H_{fGn}$ of CoP should influence the use of $\mathbf{M}$ for judgments of heaviness. It does not suggest that $H_{fGn}$ should predict the use of specifically linear or specifically quadratic components of the static moment.
<sup>‡</sup>Boldface values indicate signifance at the alpha level of 0.05.

**Fig. 4.**
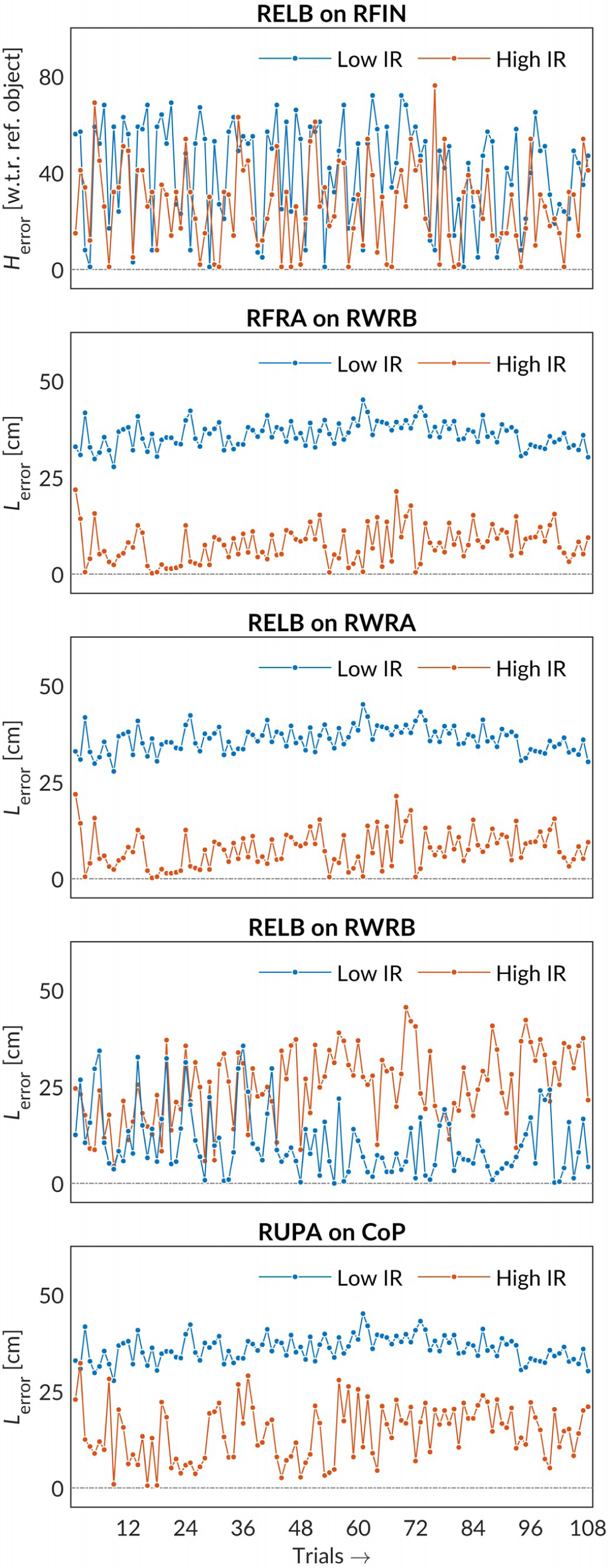
Comparisons of absolute errors in perceived heaviness, *H*_error_, and perceived length, *L*_error_, for two representative participants with low and high impulse-response (IR) values corresponding to each significant effect in Table 2. An increase in RFIN fractality due to RELB fractality reduced *H*_error_ (*p* = 0.047). An increase in CoP fractality due to RUPA fractality (*P* = 0.007), RWRB fractality due to RFRA fractality (*P* = 0.009), and RWRA fractality due to RELB fractality (*P* < 0.001) resulted decreased *L*_error_. By contrast, the flow of fractality from RELB to RWRB increased *L*_error_ (*P* = 0.004). Panels include judgments in the order the task was completed.

The flow of information through bodywide haptic perceptual system of effortful touch is bound up in each participant’s profile of dispersion of fractal fluctuations. Fig. 4 shows causal network maps showing the diffusion of fractal fluctuations — as revealed by significant IRF relationships — for the two participants who reported the least and the most accurate perceptions of length. Participants may vary in how they respond to the flux of mechanotransduction, as well as in how they coordinate a set of anatomical components to meet the task demands over time (*35*). These findings show that spatiotemporal patterns in the flow of fractality provide a snapshot into individual differences in bodywide coordination patterns underpinning perception.

## DISCUSSION

We used a network-based nonlinear approach to investigate how the bodywide dispersal and global broadcasting of local disturbances across disparate anatomical locations supports the effortful perception of object properties by manual hefting. Fluctuations in CoP and different anatomical locations showed fractality. VAR modeling revealed that the wrist and elbow spread fractality to their neighbors; as fractality in the upper arm increased, it brought down fractality among these neighbors (and the wrist); upper arm and CoP fed upon each other’s fractality. Finally, and most interestingly, the flow of perceptual information — as reflected by the accuracy of perceived heaviness and length — bound up in each participant’s profile of dispersion of fractal fluctuation. These patterns in the waxing and waning of fluctuations across disparate anatomical locations provide novel insights into how the high-dimensional flux of mechanotransduction is compressed into low-dimensional perceptual information specifying properties of hefted occluded objects.

The present results generally confirm our expectation that manually hefting an occluded object to perceive its heaviness and length should exhibit a distributed exchange of fractal fluctuations across the body. We found that CoP does have effects on the hefting arm upwards and is not just absorbing downstream fractality from the arm. Also, the sharing of fractal fluctuations across the body appears to support a greater accuracy in perceptual judgments. The role that fractal fluctuations have for predicting perceptual outcomes suggests that the participant, in effect, wears their perceptual processing on their anatomical sleeves. Quite literally, we can take fractal indicators as a way to make public the private consideration a participant makes as they come to their judgment.

More specifically, we can see three major points: one point about the lower arm, a second point about the relationship between upper arm and CoP, and a third point about the general flow of fractality that appears to support accurate perceptual judgments. First, during hefting, the lower arm (finger, wrist, and elbow) is predominantly a network of anatomical components that promotes fractality: generally, increases in fractality in any one part of the lower arm contributed to increases in fractality in other parts of the lower arm. Second, beyond this positive spread of fractality among the various parts of the lower arm, a sort of pipeline — through which fractality could flow — formed between the upper arm and CoP. Through a reciprocal relationship between the upper arm and CoP, each promoted the other’s fractality in subsequent trails, and this relationship of the upper arm with CoP promoted the ability of the upper arm to draw fractality away from the lower arm. So, our earlier results (*33*) follow from the fact that CoP fluctuations do inherit the fractality from the lower arm, but they only do so by promoting an increase in fractality at the upper arm.

The third point addressed explicitly the patterns of flows of fractal fluctuation across the body that supported greater accuracy in perceptual judgments. Regression modeling of the absolute errors in judgments suggest that the flow of fractality from the upper arm to CoP, from the wrist to the elbow, and within the wrist supported more accurate hefting by the arm, but it appears that the flow of fractality from the elbow to the wrist increased the absolute errors in perception. Hence, the most accurate judgments followed from fractal fluctuations spreading from the object through the relatively distal to proximal parts of the arm and from the upper arm to CoP.

The present findings show that fractality does not explicitly contribute to perception but instead, how fractality contributes to perception depends on where it occurs and how it flows during exploration to place the perceiving-acting participant in a heightened state of poise in which he/she becomes sufficiently open to potentiially new information. Fractality is not limited to a given point of contact between the organism and its task environment (in the present task of hefting, between the hand and the handheld object). Instead, the bodywide haptic perceptual system exhibits fractal fluctuations at apparently distinct anatomical locations, and specific patterns of flow of this fractality mediate the flow of perceptual information under the anatomical constraints of motor connectivities. While the patterns of afferent activity due to the organism-environment interaction may be ultimately integrated within the central nervous system, the perception-action system bears indicators of this process of integration. This perspective has now been successfully embraced for a few decades by the perception-action perspective of ecological psychology that views cognition as concretely embodied in performance (*43*–*47*). The present study provides glimmers of this embodied sort of cognition by showing the flow of information in the waxing and waning of fractal fluctuations across disparate anatomical locations of the body.

The standard depiction of perception is traditionally, and not surprisingly, restricted to the neural network. Mechanoreceptor activity — specifying the states of individual joint(s), muscles, tendons, and ligaments — flow to spinal neurons and then to the brain by non-interacting linear pathways. Unfortunately, such depictions fail to address the challenge of implementing afferent activity at the level of coordination and identifying when and how spatially and temporally distinct signals organize so as to inform about the states of the whole body, segments of the body, states of objects attached to the body, and how these may be engaged. Fortunately, the “ultrafast” propagations of mechanical perturbations across vast distances within biological systems have prompted physiologists and movement scientists to coin the term “preflex” to indicate a rapid, apparently motoric response that is based on mechanical tensions rather than on neural transmissions (*48, 49*). By capitalizing on the self-similar and scale-free, fractal organization of the biophysical substrate of the bodywide tensegrity (*17, 18*), preflexes constitute a means of simplifying the degrees of freedom problem which haunts the spatiotemporal organization of afferent activity.

Fractality in fluctuations at a given anatomical location implies that regardless of its size, any given event (i.e., a postural wobble) in the recorded time series influences, even if the influence is infinitesimally small in magnitude, on all subsequent events and, in like fashion, is influenced by all past events. And the specific dependence of this long-term memory on the frequency of measurement defines the fractal scaling exponent. The long-term memory in the fluctuations of the process of hefting and the scaling relation common to these fluctuations provide a window into the concinnity of afferent activity at the level of coordination. The changes in the biophysical substrate of effortful touch brought about by the changes in mechanical flux would allow an ultra-fast propagation of information, which can, in principle, support both the regulation and coordination of exploratory dynamics when engaged with an object. As opposed to the regulation and coordination by electrochemical transduction, which is slow, localized, and context-independent, the regulation and coordination brought about by the rapid propagation of mechanical flux in the bodywide, vast and complex network of connective tissues and extracellular matrix would be faster, entail local-to-global and global-to-local interactions, and be context-sensitive (*24*).

The proposal that the flow of fractality facilitates exploration is founded in the statistical relationship of fractality to diffusion. Fractal fluctuations reflect a perfect compromise between overly constrained exploration (i.e., uncorrelated fluctuations) and overly ballistic exploration (i.e., persistent fluctuations). Even in the brain, fractality is greatest in networks of integrate-and-fire stochastic spiking neurons with a mid-range of neuronal plasticity, versus extremely high or low levels of plasticity (*50, 51*). Whereas overly constrained exploration would reflect an absence of impulse-response relationships, overly ballistic exploration would reflect excessive impulse-response constrained within a narrow range of directions. The rather heterogeneous flow of fractality observed in the present study shows that during effortful touch, the body is fully poised to allow the flow of perceptual information in specific directions, reflecting how disparate anatomical components may compensate for each other based on task constraints.

The present findings, specifically the effects of IRF values on the accuracy of perceived object properties, run the risk of seeming to imply that “the stronger the fractality, or the flow of fractality, the better the perception.” We would caution against the temptation to draw any such conclusion. Instead, we would propose that stronger fractality, or the flow of fractality, places the body in a heightened state of poise, thus enabling greater access to novel information. Fractality, or more generally, long-term memory of variability, can be plainly at odds with accurately perceiving, as, for instance, the flow of fractality from the elbow to the wrist reduced the accuracy of perception of length. Previously, it has been reported that experimentally providing feedback to participants freely tapping a finger at regular, 1-s intervals increases performance at the expense of fractality in fluctuations in intertap-interval series (*52, 53*).

Perceiving an intended property of an occluded object entails a certain level of uncertainty, as each attempt at hefting an object requires a novel search. In the present experiment, even if the participants may have developed over several trials some heuristic for arriving at judgments of heaviness and length, it cannot be denied that the perceptual system must still be flexibly poised to be responsive to the randomized presentation of experimental objects. Fractal fluctuations appear to provide a common currency for the flow of information, which is not surprising as fractals provide the most efficient known way of compressing high-dimensional flux of physiological activity. Fractal fluctuations have already been shown to provide for the flexibility in neuronal activity needed by the CNS to anticipate novel structures in perceptual learning (*54*), and the present work extends the role of fractality and the flow of fractality across disparate anatomical locations of the body. Future work could investigate the general principles governing the flow of fractality and its relationship to specific goal-directed tasks (i.e., perception of heaviness versus length versus shape), as fractal fluctuations, and more generally, patterns of exploratory procedures, are strongly linked with the perceptual intent of the perceiver (*32*).

In summary, the present findings support the ecological perspective that the bodywide haptic perceptual system of effortful touch shows four defining characteristics: (1) Functionality: the components self-organize for stabilizing the task performance. (2) Flexibility: perception is not strictly dependent on specific aspects of anatomy. (3) Compensatory; disparate components reciprocally compensate for fluctuations in the environment and within the components themselves. (4) Context-sensitivity: the role of the coordinative structure as a whole or any individual component changes depending on task constraints (*55*).

## CONCLUSIONS

Despite a long history of research pointing to the importance of fractal fluctuations in physiology (*56, 57*), questions about how to link specific fractal evidence in different observables across the body remain unanswered. Specifically, it has remained unclear how fractal fluctuations might interact across the body and how those interactions might support the coordination of goal-directed behaviors. The present study was motivated by the idea that identifying the causal network structure of fractal fluctuations in the bodywide coordination may be a fruitful way of understanding the haptic perceptual capabilities of effortful touch at the level of the underlying coordination. It provides a compeling evidence that a complex interplay of fractality in mechanical fluctuations at disparate anatomical locations of the body support perception via effortful touch. The present study is a significant step towards the solution of a fundamental problem in human perception: how is afferent activity diffused throughout the body unified as an instance of conscious perceptual experience? Fractal fluctuations are a promising candidate for engaging disparate components of the bodywide tensegrity into a coherent activity and provide a strategy for the local-to-global and global-to-local exchange of information, thus ensuring the completeness of a transformation from diffused afferent activity into conscious perceptual experience. The flow of fractality in perception-action tasks could be studied using causal network analysis as a common framework, potentially providing novel insights and interventions into conditions such as developmental coordination disorder (DCD) and attention-deficit hyper disorder (ADHD) that narrow the spectrum of individuals’ psychomotor complexity (*28, 58*).

## MATERIALS AND METHODS

### Participants

Eight adult men and seven adult women [*Mean* (±1*SD*) = 23.4 (3.4) years, all right-handed] without any self-reported neurological or sensorimotor disorder voluntarily participated in the present study. Each participant provided verbal and written consent after being informed about the purposes of the study, the procedures, and the potential risks and benefits of participation, in compliance with the Declaration of Helsinki. The Institutional Review Board (IRB) at the University of Georgia (Athens, GA) approved the present study.

### Experimental objects

Each participant hefted six experimental objects, each consisting of an oak, hollow aluminum, or solid aluminum dowel (diameter = 1.2 cm, length = 75.0 cm; mass = 68 g, 109 g, and 266 g, respectively) weighted by either 4 or 12 stacked steel rings attached at 20.0 or 60.0 cm, respectively (inner diameter = 1.4 cm, outer diameter = 3.4 cm, thickness = 0.8 cm and 2.4 cm, respectively; mass = 56 g and 168 g, respectively) (Table 1 and Fig. 1A). The dowels were weighted such that the resulting six objects systematically differed in their mass, *m* (Object 1 > Object 2, Object 3 > Object 4, Object 5 > Object 6), the static moment, **M** (Object 1 = Object 2 = **M**_S_ < Object 3 = Object 4 = **M**_M_ < Object 5 = Object 6 = **M**_L_), and the moment of inertia, *I*_1_ and *I*_3_, reflecting the resistance of the object to rotation about the longitudinal axis (*I*_1_ values: Object 1, Object 2, Object 3 < Object 4, Object 5 < Object 6). A cotton tape of negligible mass was enfolded on each dowel to prevent the cutaneous perception of its composition.

### Experimental setup and procedure

After being blindfolded, each participant stood with each foot on separate force plates (60×40 cm; Bertec Inc., Columbus, OH), hefted each object, and reported judgments of heaviness and length (Fig. 1B). The participant was asked to constrain his/her wrist motion about 10° ulnar deviation, the neutral position, or 10° radial deviation (Fig. 1C). A custom setup consisting of two tripods supported the object such that the object was aligned parallel to the participant’s wrist. The inclusion of the different wrist angles allowed us to investigate the effects the postural constraints on hefting and wielding on perceptual judgments of heaviness and length. In a static condition, the participant lifted and held each object static. In two dynamic conditions, instead of freely hefting the objects — which has been traditionally done in dynamic or effortful touch tasks — in the dynamic condition, the participant lifted and wielded each object synchronously with metronome beats at 2 Hz or 3 Hz, which added additional constraints on perceptual exploration. The participant was instructed to minimize the motion of the torso and upper hand, and the amplitude of wielding movements.

### Experimental setup and procedure

To track the motion of the hefted object and that of the participant’s body in 3D, we attached using double-sided adhesive tape three reflective markers (diameter = 9.5 mm) on each experimental object at 30, 45, and 60 cm from the object’s proximal end and nine reflective markers on the participant’s body (Supplementary Table S1 and Fig. 2A). We tracked the 3D motion of of each reflective marker at 100 Hz using an eight-camera Qualisys motion tracking system (Qualisys Inc., Boston, MA) as a participant hefted an object.

Each participant completed a total of 108 trials (3 Wrist angles × 3 Wrist angular kinematics × 6 Objects × 2 Trials/Object) in a 90–105-min session. A nested, pseudo-randomized block design was used, the factors of Wrist angular kinematics (Static, 2 Hz dynamic, and 3 Hz dynamic) being nested within the factors of Wrist angle (Radial, Neutral, and Ulnar). The order of the 12 trials (6 Objects × 2 Trials/Object) was pseudo-randomized for each block.

Before the first and after every six trials, each participant hefted a reference object that was arbitrarily attributed to a heaviness value of 100 units. Each participant was instructed to assign heaviness values proportionally higher and lower than 100 to an object perceived heavier and lighter, respectively, than the reference object (e.g., 200 to an object perceived twice as heavy and 50 to an object perceived half as heavy). In each trial, after a ‘lift’ signal, the participant lifted the object by about 5 cm and held it static or wielded it at 2 Hz or 3 Hz. After 5 s, following a ‘stop’ signal, the participant placed the object back and reported (a) perceived heaviness (no units) and (b) perceived length by adjusting the position of a marker on a string-pulley assembly. The experimenter noted the perceived length (cm) from a meter-scale attached to the base of the string-pulley assembly and occluded from the participant.

### Data processing

#### CoP planar Euclidean displacement (PED) series

The output of force plates was downsampled by 1/20 (i.e., from 2000 Hz to 100 Hz) to match the sampling rates of kinematic trajectories of reflective markers and the ground reaction forces. The ground reaction forces recorded on each trial yielded a two-dimensional center of pressure (CoP) time series, with each dimension describing the position of the CoP along the participant’s medial: lateral and anterior: posterior axes. Recording on each trial over 5 s yielded a two-dimensional CoP time series of 500 samples and thus the corresponding CoP displacement time series consisting of 499 samples. Finally, a one-dimensional CoP planar Euclidean displacement (PED) series was obtained for each downsampled CoP time series, describing CoP displacement along the transverse plane of the body (Fig. 2B).

#### Body sway displacement series

Motion tracking of each reflective marker attached to the body and the experimental objects (*N* = 12) yielded a three-dimensional kinematic time series, with each dimension describing the position of the marker along the participant’s medial: lateral, anterior: posterior, and superior: inferior axes. Recording on each trial over 5 s yielded a three-dimensional sway time series of 500 samples and thus the corresponding time series of marker displacement consisting of 499 samples. Finally, a one-dimensional spatial Euclidean displacement (SED) series was obtained for each marker describing the displacement of that marker in 3D (Fig. 2B).

#### Detrended fluctuation analysis

We used detrended fluctuation analysis (DFA) to compute the Hurst exponent, *H*, describing the strength of temporal correlations in the PED series. DFA was first developed to estimate the strength of temporal correlations in a given time series (*37, 38*). The DFA proceeds by finding the first-order integration of a time series *x* (*t*) with *N* samples to compute the cumulative sums of difference scores to produce the new time series:

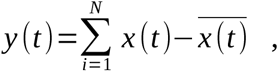

where 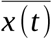 is the grand mean of the time series. Next, a linear trend *y*_*n*_ (*t*) is fit to nonoverlapping n-length bin of *y* (*t*) and the root mean square (RMS; i.e., averaging the residuals) over each fit is computed. RMS over each bin size constitutes a fluctuation function *f* (*N*):

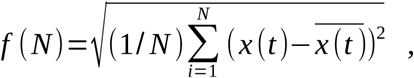

for *n*<*N*/ 4. On standard scales, *f* (*N*) is a power law:

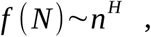

where *H* is the scaling exponent. The closer *H* is to 1, the stronger the temporal correlations are. *H* is estimated by logarithmically scaling the previous equation:

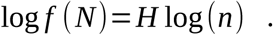

Hence, the slope of fluctuation functions in log-log plots represents *H*. It is important to note that temporal correlations can be present in both a time series and its first-order derivative. The original time series are often classified as fractional Brownian motions (*fBm*), wherein the first-order derivative of *fBm* is fractional Gaussian noise (*fGn*). Accordingly, the scaling exponents of a trajectory and its first-order derivative are denoted *H*_*fBm*_ and *H*_*fGn*_, respectively.

We obtained DFA estimates for the original version (i.e., unshuffled) and a shuffled version (i.e., a version with the temporal information destroyed) of each CoP PED series, as well as of each marker SED series, over each of the following bin sizes: 4, 8, 12,… 128 (Fig. 2C). Exploration at the level of individual trials indicated inflection points in fluctuation functions, specifically at larger timescales. To test for this possibility, we also obtained DFA estimates for the original version and a shuffled version of each CoP PED series, as well as of each marker SED series, over half of the scaling region: 4, 8, 12,… 64.

#### Vector autoregression analysis

Vector autoregression (VAR) is a technique for modeling stochastic processes to capture the linear interdependencies among multiple time series. The evolution of each entered variable is described by an equation based on its own lagged value and that of each other variable, along with an error term. As compared to structural models that require prior knowledge of the factors influencing a variable, the only prior knowledge required for VAR modeling is a list of variables that can be hypothesized to affect each other intertemporally.

VAR can produce a system of *m* regression equations predicting each variable as a function of lagged values of themselves and of each other. In the simplest case of *m*=2, with a pair of time series *f* (*t*) and *g*(*t*) definable at each value of time *t* =1 to *t* =*N*, where *N* is the length of the time series, a VAR model would have the following structure:

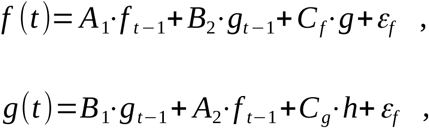

where *A* _*j*_ and *B* _*j*_ are the coefficients quantifying the effects of the previous values of *f* and *g*, respectively, with *j* indexing the variable to which these previous values contribute and with error terms *ε*_*f*_ and *ε*_*g*_ (*59*). The above equations describe a 1-lag VAR, that is, each *f* and *g* is described in terms of values up to 1 value preceding the predicted values. VAR models can include exogenous variables, such as the factors of experimental design, which stand outside the mutual relationship among the variables internal to the system. In the above example, the time series *h*(*t*) can induce changes in *f* (*t*) or *g*(*t*), but changes in neither *f* (*t*) or *g*(*t*) can induce changes in *h*(*t*). *h* is an exogenous variable, and *C*_*f*_ and *C*_*g*_ are coefficients indicating the effect of *h*(*t*) on *f* (*t*) and *g*(*t*), respectively. Endogenous variables are variables internal to the system (i.e., *f* (*t*) or *g*(*t*)), which may respond to and induce changes in other endogenous variables. For the purposes of the present analysis, the fractal scaling exponent corresponding to each of the 13 anatomical locations (CoP and the 12 reflective markers) served as an endogenous variable (Fig. 2D).

VAR models provide forecasts of the effects of endogenous variables into the future through impulse-response functions (IRFs). Whereas standard regression evaluates the relationship between *f* (*t*) and *g*(*t*), IRFs can evaluate relationships between *f* (*t*) and *g*(*t* + *τ*), or between *g*(*t*) and *f* (*t* + *τ*), where *τ* is a whole number. First, orthogonalizing the regression equations and, second, inducing an ‘impulse’ to the system of regression equations by adding 1 standard error to any single variable, propogates responses across variables. The plot of an IRF describes the changes in predicted later values of one time series due to the impulse from another time series (*59, 60*). It is customary to fit the least number of lags that leave independently and identically distributed residuals. VAR modeling does not require as much knowledge about the forces influencing a variable; the only prior knowledge required is a list of variables which can be hypothesized to affect each other intertemporally, thus allowing us to explore causal relationships after addressing minimal short-lag relationships (*61*).

#### Statistical analysis

The goal was to understand how the fractal scaling exponents (DFAs) for the 13 locations (corresponding to CoP and the 12 reflective markers attached to various body parts) differed in the following ways: (1) the DFA at each location may differ in its average effect as an impulse variable on the DFAs at all locations (the global impulse effect). (2) The DFA at each location may differ in its response to the DFAs at all locations (the global response effect). (3) Each pairwise relationship between the DFAs at the 13 locations may show specifically different impulse-response relationships than for the first two global cases (the specific pairwise impulse-response effect).

All impulse-response relationships indicating the subsequent effects of increases in the DFAs were submitted to a full-factorial regression model (*62*) using the “nlme” package for RStudio (*63*). A full-factorial regression model of Impulse × Response × Trial was used, with Impulse and Response serving as class variables indicating the locations of the impulse variables and the responding variables, respectively. The regression utilized orthogonal linear, quadratic, and cubic polynomials to model the impulse-response relationships. The Impulse terms in this full-factorial design allowed estimating the global effect of the prior increase in the DFA of each location on the intercept and the linear, quadratic, and cubic components of all impulse-response relationships. The Response terms in this full-factorial design allowed estimating the effect of the subsequent increase in the DFA at each location on the intercept and the linear, quadratic, and cubic components of all impulse-response relationships. Thus, the Impulse and Response effects would portray the tendency for the DFA at specific locations to influence or to be influenced according to different third-order polynomial responses over subsequent trials. The Impulse × Response terms would highlight significant differences of specific pairs of impulse and response variables for which the impulse-response relationship deviated from the global patterns.

Finally, we modeled the accuracy of perceptual judgments, encoded by the unsigned error in judgments: absolute(*H*/*L*_*perceived*_ - *H*/*L*_actual_). For calculating the signed error in perceived length, we subtracted the actual length (i.e., 75 cm) from perceived length. Because perceived heaviness followed a proportion relative to a reference object of 109-gm, we calculated this judgment as the percentage of the [theoretically] accurate percentage value based on each object’s actual mass. For instance, if a participant perceived Object 2 to have a length of 62.5 cm and heaviness 120 relative to 100 of the referenced object, then they would have signed error in perceived length, *L*_error_ = 62.5 − 75.0 = − 12.5 and signed error in perceived heaviness, *H*_error_ = 100×((120×109)/100)/236 = 55.42. Next, for calculating the unsigned error, we calculated the absolute value of error in perceived length, and the absolute value of 100 less than the percentage value corresponding to *H*_perceived_. Accordingly, for perceptions of the same object, the unsigned error in perceived length would be 12.5, and the unsigned error in perceived heaviness would be the absolute value of 55.34 – 100 = 44.66. We rounded the percentage error values to the nearest integer.

Perceived heaviness was a nonlinear dependent measure, given the instruction to report heaviness in terms of ratios to the reference object (e.g., 200 to an object perceived twice as heavy and 50 to an object perceived half as heavy). So, it is evident that the dependent measure is just as skewed as that multiplicative definition should indicate. Thus, rather than submitting the data to two steps of 1) a log transformation, and 2) linear regression (i.e., use logistic regression), to accommodate this skew, we used the generalized linear model (GLM) of Poisson regression, which is much like logistic regression but uses a log link instead of a logit link function. By contrast, perceived length was explicitly linear as we defined it. Accordingly, we used the GLM of Poisson regression using “lme4” package for Rstudio (*64*) to examine variation in unsigned error in perceived heaviness; and linear mixed-effect (LME) models using the “nlme” package for RStudio (*63*), to examine variation in unsigned error in perceived length.

Predictors included Trial order, Wrist angle, Wrist angular kinematics, Object’s static moment, logarithmic of object’s moments of inertia (Log*I*_1_ and Log*I*_3_), fractal scaling exponent *H*_fGn_ at CoP, and the IRF values forcasting the response to impulse in the first subsequent trial for the following IR relationships: CoP on RUPA, RUPA on CoP, RFRA on RFIN, RFRA on RFRB, RELB on RFIN, RELB on RWRA, and RELB on RWRB. Wherever possible, we fit the effects of both the static moment and the moments of inertia, respecting the fact that these different aspects of the mass distribution can play a role in perceived heaviness and perceived length (*65, 66*), but this policy worked best in the model for perceived heaviness. The ordinal encoding of the static moment (i.e., **M**_S_, **M**_M_, and **M**_L_) required that we fit orthogonal polynomials to allow for the possibility of both linear and quadratic effects of this variable. When modeling did not support the inclusion of all object properties (mass, the static moment, and the moment of inertia, we resorted to modeling length perception as a function of the moment of inertia to the exclusion of other properties. Crucially, perception hinges on the relevance of interactions between H_fGn_ at CoP and object parameters (*33*), and thus we included this interaction as well.

## SUPPLEMENTARY MATERIALS

**Table S1.** Location of the reflective markers attached to each experimental object and the participant’s body

|  | **Marker** | **Location** |
| --- | --- | --- |
| Experimental object | OBJP | Tip of the object |
|  | OBJD | 30 cm from the distal end |
|  | OBJP | 30 cm from the proximal end |
| Participant's body | RFIN | Just below the middle knuckle on the right hand |
|  | RWRA | Extended from the thumb side using a wrist bar |
|  | RWRB | Extended from the little finger side using a wrist bar |
|  | RFRM | On the outside of the lower arm |
|  | RELB | On the bony prominence on the outside of the elbow joint |
|  | RUPA | Outside of the upper arm |
|  | RSHO | On the bony prominence on top of the right shoulder |
|  | CLAV | Top of the breast bone |
|  | STRN | Base of the breast bone |

**Table S2.** *Mean* (±1*SEM*) values of *H*_fGn_ yielded by DFA for the original and a shuffled version of each CoP PED and marker SED series, and coefficients of paired samples *t*-tests comparing the two

| **Location** | **$H_{fGn}$ (unshuffled)** | **$H_{fGn}$ (shuffled)** | **$t_{2,1619}$** | **$P^\dagger$** |
| --- | --- | --- | --- | --- |
| OBJT | 1.14 (0.0083) | 0.51 (0.0016) | 73.46 | < **0.000** |
| OBJD | 1.15 (0.0083) | 0.51 (0.0015) | 76.20 | < **0.000** |
| OBJP | 1.74 (0.0078) | 0.51 (0.0021) | 80.24 | < **0.000** |
| RFIN | 1.89 (0.0059) | 0.52 (0.0018) | 109.54 | < **0.000** |
| RWRA | 1.19 (0.0059) | 0.51 (0.0021) | 112.59 | < **0.000** |
| RWRB | 1.15 (0.0059) | 0.51 (0.0015) | 104.11 | < **0.000** |
| RFRM | 1.10 (0.0048) | 0.51 (0.0014) | 118.06 | < **0.000** |
| RELB | 1.06 (0.0044) | 0.51 (0.0013) | 118.08 | < **0.000** |
| RUPA | 0.97 (0.0048) | 0.52 (0.0024) | 79.80 | < **0.000** |
| RSHO | 0.97 (0.0048) | 0.51 (0.0025) | 83.69 | < **0.000** |
| CLAV | 0.89 (0.0049) | 0.51 (0.0030) | 69.68 | < **0.000** |
| STRN | 0.89 (0.0045) | 0.51 (0.0027) | 72.46 | < **0.000** |
| CoP | 0.57 (0.0018) | 0.51 (0.0013) | 25.57 | < **0.000** |

<sup>†</sup>Boldface values indicate signifiacne at the two-tailed alpha level of 0.05.

**Table S3.** *Mean* (±1*SEM*) values of *H*_fGn_ yielded by DFA for the original and a shuffled version of each CoP PED and marker SED series for a shorter, half of the scaling region, and coefficients of paired samples *t*-tests comparing the two

| **Location** | **$H_{fGn}$ (unshuffled)** | **$H_{fGn}$ (shuffled)** | **$t_{2,1619}$** | **$P^{\dagger}$** |
| --- | --- | --- | --- | --- |
| OBJP | 1.24 (0.0085) | 0.53 (0.0017) | 81.32 | < **0.000** |
| OBJD | 1.25 (0.0082) | 0.53 (0.0016) | 84.18 | < **0.000** |
| OBJP | 1.25 (0.0073) | 0.53 (0.0014) | 94.56 | < **0.000** |
| RFIN | 1.21 (0.0053) | 0.53 (0.0013) | 123.04 | < **0.000** |
| RWRA | 1.21 (0.0054) | 0.54 (0.0029) | 109.88 | < **0.000** |
| RWRB | 1.61 (0.0053) | 0.53 (0.0014) | 114.16 | < **0.000** |
| RFRM | 1.10 (0.0042) | 0.53 (0.0012) | 130.91 | < **0.000** |
| RELB | 1.08 (0.0039) | 0.53 (0.0012) | 135.41 | < **0.000** |
| RUPA | 0.97 (0.0044) | 0.53 (0.0039) | 75.09 | < **0.000** |
| RSHO | 1.00 (0.0045) | 0.54 (0.0045) | 72.30 | < **0.000** |
| CLAV | 0.91 (0.0049) | 0.53 (0.0031) | 60.55 | < **0.000** |
| STRN | 0.90 (0.0046) | 0.54 (0.0031) | 61.43 | < **0.000** |
| CoP | 0.60 (0.0014) | 0.53 (0.0011) | 36.44 | < **0.000** |

<sup>†</sup>Boldface values indicate signifiacne at the two-tailed alpha level of 0.05.

**Table S4. Complete output of the full-factorial regression model of Impulse × Response × Trial, with Impulse and Response serving as class variables indicating the locations of the impulse variables and the responding variables, respectively**

## Author contributions

M.M. conceived and designed research; M.M. performed experiments; M.M., N.S.C., and D.G.K-S. analyzed data; M.M. and D.G.K-S. interpreted results of experiments; M.M. prepared figures; M.M. and D.G.K-S. drafted manuscript; M.M., N.S.C., and D.G.K-S. edited and revised manuscript; M.M., N.S.C., and D.G.K-S. approved final version of manuscript.

## Data and materials availability

All data needed to evaluate the conclusions in the paper are present in the paper and/or the Supplementary Materials. Additional data related to this paper may be requested from the authors.

## REFERENCES AND NOTES

1. G. Burton, M. T. Turvey, H. Y. Solomon, Can shape be perceived by dynamic touch? Percept. Psychophys. 48, 477–487 (1990).

2. C. Carello, P. Fitzpatrick, I. Flascher, M. T. Turvey, Inertial eigenvalues, rod density, and rod diameter in length perception by dynamic touch. Percept. Psychophys. 60, 89–100 (1998).

3. C. Carello, M. T. Turvey, in Touch, Representation and Blindness, M. A. Heller, Ed. (Oxford University Press, New York, NY, 2000), pp. 27–66.

4. M. T. Turvey, G. Burton, E. L. Amazeen, M. Butwill, C. Carello, Perceiving the width and height of a hand-held object by dynamic touch. J. Exp. Psychol. Hum. Percept. Perform. 24, 35–48 (1998).

5. M. T. Turvey, G. Burton, C. C. Pagano, H. Y. Solomon, S. Runeson, Role of the inertia tensor in perceiving object orientation by dynamic touch. J. Exp. Psychol. Hum. Percept. Perform. 18, 714–727 (1992).

6. C. C. Pagano, P. Fitzpatrick, M. T. Turvey, Tensorial basis to the constancy of perceived object extent over variations of dynamic touch. Percept. Psychophys. 54, 43–54 (1993).

7. G. Burton, M. T. Turvey, Attentionally splitting the mass distribution of hand-held rods. Percept. Psychophys. 50, 129–140 (1991).

8. C. C. Pagano, M. T. Turvey, Eigenvectors of the inertia tensor and perceiving the orientation of a hand-held object by dynamic touch. Percept. Psychophys. 52, 617–624 (1992).

9. M. T. Turvey, C. Carello, Obtaining information by dynamic (effortful) touching. Philos. Trans. R. Soc. London B Biol. Sci. 366, 3123–3132 (2011).

10. J. B. Wagman, A. Hajnal, Task specificity and anatomical independence in perception of properties by means of a wielded object. J. Exp. Psychol. Hum. Percept. Perform. 40, 2372–2391 (2014).

11. J. B. Wagman, A. Hajnal, Getting off on the right (or left) foot: Perceiving by means of a rod attached to the preferred or non-preferred foot. Exp. Brain Res. 232, 3591–3599 (2014).

12. J. B. Wagman, M. D. Langley, T. Higuchi, Turning perception on its head: Cephalic perception of whole and partial length of a wielded object. Exp. Brain Res. 235, 153–167 (2017).

13. D. G. Stephen, A. Hajnal, Transfer of calibration between hand and foot: Functional equivalence and fractal fluctuations. Attention, Perception, Psychophys. 73, 1302–1328 (2011).

14. A. Hajnal, S. Fonseca, S. Harrison, J. M. Kinsella-Shaw, C. Carello, Comparison of dynamic (effortful) touch by hand and foot. J. Mot. Behav. 39, 82–88 (2007).

15. Z. Palatinus, C. Carello, M. T. Turvey, Principles of part–whole selective perception by dynamic touch extend to the torso. J. Mot. Behav. 43, 87–93 (2011).

16. A. Hajnal et al., Haptic selective attention by foot and by hand. Neurosci. Lett. 419, 5–9 (2007).

17. M. T. Turvey, S. T. Fonseca, The medium of haptic perception: A tensegrity hypothesis. J. Mot. Behav. 46, 143–187 (2014).

18. P. A. Cabe, All perception engages the tensegrity-based haptic medium. Ecol. Psychol., 1–13 (2018).

19. D. E. Ingber, Cellular mechanotransduction: Putting all the pieces together again. FASEB J. 20, 811–827 (2006).

20. D. E. Ingber, From cellular mechanotransduction to biologically inspired engineering. Ann. Biomed. Eng. 38, 1148–1161 (2010).

21. D. G. Kelty-Stephen, Multifractal evidence of nonlinear interactions stabilizing posture for phasmids in windy conditions: A reanalysis of insect postural-sway data. PLoS One. 13, e0202367 (2018).

22. D. E. Ingber, Tensegrity-based mechanosensing from macro to micro. Prog. Biophys. Mol. Biol. 97, 163–179 (2008).

23. D. E. Ingber, Tensegrity and mechanotransduction. J. Bodyw. Mov. Ther. 12, 198–200 (2008).

24. M. T. Turvey, Action and perception at the level of synergies. Hum. Mov. Sci. 26, 657–697 (2007).

25. W. H. Warren, in Sensory-Motor Organizations and Development in Infancy and Early Childhood, B. Bloch, B. I. Bertenthal, Eds. (Springer, Dordrecht, Netherlands, 1990), pp. 23–37.

26. G. C. Van Orden, J. G. Holden, M. T. Turvey, Self-organization of cognitive performance. J. Exp. Psychol. Gen. 132, 331–350 (2003).

27. C. T. Kello, Critical branching neural networks. Psychol. Rev. 120, 230–254 (2013).

28. B. S. Avelar et al., Fractal fluctuations in exploratory movements predict differences in dynamic touch capabilities between children with Attention-Deficit Hyperactivity Disorder and typical development. PLoS One. 14, e0217200 (2019).

29. D. G. Stephen, R. Arzamarski, C. F. Michaels, The role of fractality in perceptual learning: Exploration in dynamic touch. J. Exp. Psychol. Hum. Percept. Perform. 36, 1161–1173 (2010).

30. M. Mangalam, J. D. Conners, D. G. Kelty-Stephen, T. Singh, Fractal fluctuations in muscular activity contribute to judgments of length but not heaviness via dynamic touch. Exp. Brain Res. 237, 1213–1216 (2019).

31. Z. Palatinus, J. A. Dixon, D. G. Kelty-Stephen, Fractal fluctuations in quiet standing predict the use of mechanical information for haptic perception. Ann. Biomed. Eng. 41, 1625–1634 (2013).

32. Z. Palatinus, D. G. Kelty-Stephen, J. Kinsella-Shaw, C. Carello, M. T. Turvey, Haptic perceptual intent in quiet standing affects multifractal scaling of postural fluctuations. J. Exp. Psychol. Hum. Percept. Perform. 40, 1808–1818 (2014).

33. M. Mangalam, R. Chen, T. R. McHugh, T. Singh, D. G. Kelty-Stephen, Bodywide fluctuations support manual exploration: Fractal fluctuations in posture predict perception of heaviness and length via effortful touch by the hand. Hum. Mov. Sci. 69, 102543 (2020).

34. M. Mangalam, D. G. Kelty-Stephen, Multiplicative-cascade dynamics supports whole-body coordination for perception via effortful touch. Hum. Mov. Sci. (2020).

35. D. G. Kelty-Stephen, J. A. Dixon, Interwoven fluctuations during intermodal perception: Fractality in head sway supports the use of visual feedback in haptic perceptual judgments by manual wielding. J. Exp. Psychol. Hum. Percept. Perform. 40, 2289–2309 (2014).

36. L. Kilian, H. Lütkepohl, Structural vector autoregressive analysis (Cambridge University Press, Cambridge, UK, 2017).

37. C.-K. Peng et al., Mosaic organization of DNA nucleotides. Phys. Rev. E. 49, 1685–1689 (1994).

38. C.-K. Peng, S. Havlin, H. E. Stanley, A. L. Goldberger, Quantification of scaling exponents and crossover phenomena in nonstationary heartbeat time series. Chaos An Interdiscip. J. Nonlinear Sci. 5, 82–87 (1995).

39. L. S. Liebovitch, W. Yang, Transition from persistent to antipersistent correlation in biological systems. Phys. Rev. E. 56, 4557–4566 (1997).

40. R. Baldwin, P. Krugman, Persistent trade effects of large exchange rate shocks. Q. J. Econ. 104, 635–654 (1989).

41. B. J. West, E. L. Geneston, P. Grigolini, Maximizing information exchange between complex networks. Phys. Rep. 468, 1–99 (2008).

42. B. J. West, N. Scafetta, Nonlinear dynamical model of human gait. Phys. Rev. E. 67, 51917 (2003).

43. E. J. Gibson, A. D. Pick, Ecological Approach to Perceptual Learning and Development (Oxford University Press, New York, NY, 2000).

44. J. J. Gibson, The Senses Considered as Perceptual Systems (Houghton Mifflin, Boston, MA, 1966).

45. A. Chemero, Radical Embodied Cognitive Science (MIT Press, Cambridge, MA, 2009).

46. A. Clark, N. Eilan, Sensorimotor skills and perception. Proc. Aristot. Soc. Suppl. Vol. 80, 43–88 (2006).

47. W. H. Warren, The dynamics of perception and action. Psychol. Rev. 113, 358–389 (2006).

48. A. B. Chambliss et al., The LINC-anchored actin cap connects the extracellular milieu to the nucleus for ultrafast mechanotransduction. Sci. Rep. 3, 1087 (2013).

49. Z. Jahed, H. Shams, M. R. K. Mofrad, A disulfide bond Is required for the transmission of forces through SUN-KASH complexes. Biophys. J. 109, 501–509 (2015).

50. A. A. Costa, M. J. Amon, O. Sporns, L. H. Favela, in International Workshop on Complex Networks: Complex Networks IX, S. Cornelius, K. Coronges, B. Gonçalves, R. Sinatra, A. Vespignani, Eds. (Springer International Publishing, Cham, 2018), pp. 161–171.

51. D. Aguilar-Velázquez, L. Guzmán-Vargas, Synchronization and 1/*i* signals in interacting small-world networks. Chaos, Solitons & Fractals. 104, 418–425 (2017).

52. N. Kuznetsov, S. Wallot, Effects of accuracy feedback on fractal characteristics of time estimation. Front. Integr. Neurosci. 5, 62 (2011).

53. A. Eke et al., Pitfalls in fractal time series analysis: fMRI BOLD as an exemplary case. Front. Physiol. 3 (2012), p. 417.

54. E. Bieberich, Recurrent fractal neural networks: A strategy for the exchange of local and global information processing in the brain. Biosystems. 66, 145–164 (2002).

55. B. J. Thomas, M. A. Riley, J. B. Wagman, in Perception as Information Detection: Reflections on Gibson’s Ecological Approach to Visual Perception, J. B. Wagman, J. J. C. Blau, Eds. (Routledge, New York, NY, 2019), pp. 237–252.

56. R. W. Glenny, H. T. Robertson, S. Yamashiro, J. B. Bassingthwaighte, Applications of fractal analysis to physiology. J. Appl. Physiol. 70, 2351–2367 (1991).

57. D. A. Beard, J. B. Bassingthwaighte, The fractal nature of myocardial blood flow emerges from a whole-organ model of arterial network. J. Vasc. Res. 37, 282–296 (2000).

58. J. M. Ocarino et al., Dynamic touch is affected in children with cerebral palsy. Hum. Mov. Sci. 33, 85–96 (2014).

59. H. Lutkepohl, New Introduction to Multiple Time Series Analysis (Springer, New York, NY, 2007).

60. A. Hatemi-J, Multivariate tests for autocorrelation in the stable and unstable VAR models. Econ. Model. 21, 661–683 (2004).

61. C. A. Sims, Macroeconomics and reality. Econometrica. 48, 1–48 (1980).

62. J. D. Singer, J. B. Willett, Applied Longitudinal Analysis: Modeling Change and Event Occurrence (Oxford University Press, New York, NY, 2003).

63. J. Pinheiro, D. Bates, S. DebRoy, D. Sarkar, R. C. Team, nlme: Linear and nonlinear mixed effects models. R Packag. version 3.1-137 (2018) (available at https://cran.r-project.org/package=nlme).

64. D. Bates, D. Sarkar, M. Bates, L. Matrix, The lme4 package (2007).

65. I. Kingma, P. J. Beek, J. H. van Dieën, The inertia tensor versus static moment and mass in perceiving length and heaviness of hand-wielded rods. J. Exp. Psychol. Hum. Percept. Perform. 28, 180–191 (2002).

66. I. Kingma, R. van de Langenberg, P. J. Beek, Which mechanical invariants are associated with the perception of length and heaviness of a nonvisible handheld rod? Testing the inertia tensor hypothesis. J. Exp. Psychol. Hum. Percept. Perform. 30, 346–354 (2004).

